# Xolotl: An Intuitive and Approachable Neuron and Network Simulator for Research and Teaching

**DOI:** 10.1101/394973

**Authors:** Srinivas Gorur-Shandilya, Alec Hoyland, Eve Marder

## Abstract

Conductance-based models of neurons are used extensively in computational neuroscience. Working with these models can be challenging due to their high dimensionality and large number of parameters. Here, we present a neuron and network simulator built on a novel automatic type system that binds object-oriented code written in C++ to objects in MATLAB. Our approach builds on the tradition of uniting the speed of languages like C++ with the ease-of-use and feature-set of scientific programming languages like MATLAB. Xolotl allows for the creation and manipulation of hierarchical models with components that are named and searchable, permitting intuitive high-level programmatic control over all parts of the model. The simulator’s architecture allows for the interactive manipulation of any parameter in any model, and for visualizing the effects of changing that parameter immediately. Xolotl is fully featured with hundreds of ion channel models from the electrophysiological literature, and can be extended to include arbitrary conductances, synapses, and mechanisms. Several core features like bookmarking of parameters and automatic hashing of source code facilitate reproducible and auditable research. Its ease of use and rich visualization capabilities make it an attractive option in teaching environments. Finally, xolotl is written in a modular fashion, includes detailed tutorials and worked examples, and is freely available at https://github.com/sg-s/xolotl, enabling seamless integration into the workflows of other researchers.

## 1 INTRODUCTION

Nervous systems process and transmit information using electrically excitable membranes. Conductancebased models are a powerful biophysical simplification of an electrically excitable compartment in a neuron (Hodgkin and Huxley 1952a). Studies based on the Hodgkin-Huxley formalism now contribute significantly to mainstream research in some circuits (Marder and Abbott 1995; Prinz 2010; Prinz 2006). These models provide an approachable framework for understanding many salient principles of electrophysiology, since they explicitly model cell membranes and ion channels as electrical components in a circuit. However, several challenges remain in working with biophysically-detailed conductance-based neuron models. First, these models can be high-dimensional with many nonlinear differential equations, each with several parameters. Second, many or all equations in these models can be strongly coupled through dynamical variables like the membrane potential. In multi-compartment models of spatially extended neurons, membrane potentials in every compartment can be different, and are coupled to each other. Finally, the choice of programming language used to implement the model imposes tradeoffs in designing and using neuron and network simulators: simulators written in languages like C++ or FORTRAN can integrate equations quickly, but often lack the ease-of-use and interoperability of those written in scientific programming languages like Python, Julia, or MATLAB (Mathworks).

Two major approaches have dominated the design of neuron simulators. One approach is to write the simulator in a fast compiled language like C and allow for the construction and simulation of neuron models using object-oriented paradigms. This approach has been implemented in NEURON (Hines and Carnevale 1997). Simulators designed in this way tend to perform fast computations with little overhead, but suffer from a steep learning curve. Wrapping these simulators in a more approachable language like Python or using graphical user interfaces (GUIs) mitigates these drawbacks (Hines, Davison, and Muller 2009; Gratiy et al. 2018) at the cost of obfuscating the underlying algorithms and parameters (Brette et al. 2007; Hines, Davison, and Muller 2009). In contrast, simulators designed from the ground up in popular scientific computing languages can be easier to use and benefit from interoperability with other commonly-used tools. Simulators like DynaSim (Sherfey et al. 2018), ANNarchy (Vitay, Dinkelbach, and Hamker 2015), BRIAN (Stimberg et al. 2013), morphforge (Hull and Willshaw 2014), and PyNN (Davison et al. 2009) allow the user to specify models with strings of equations or components that are constructed using a special syntax, that can then be translated into a faster implementation language such as C or C++ (Stimberg et al. 2014a). This approach permits considerable flexibility for simulating systems of differential equations. Because models need to be translated between the two languages, the hierarchical nature of neuron models is not naturally encapsulated by these tools, and the syntax can be verbose. Neither approach facilitates the creation of tools that simultaneously maintain efficiency, ease-of-use, and clarity.

To overcome these design limitations, we have developed a novel automatic type system, that we call cpplab, which binds MATLAB code to classes specified in C++ header files. This architecture automatically creates objects in MATLAB that represent the underlying object-oriented C++ code, allowing the symbolic manipulation of C++ objects in the MATLAB interface. In this paper, we introduce xolotl, an implementation of the cpplab system specialized in integrating conductance-based neuron and network models. Models can be easily constructed from components of different types in a few lines of MATLAB code using a hierarchical and intuitive syntax. Since models in the MATLAB workspace are automatically linked to models in the C++ implementation, configuring these objects in MATLAB transparently configures the underlying C++ objects. Xolotl comes packaged with hundreds of components that can be used to assemble cells and networks; has built-in visualization functions to inspect voltage time traces and activation functions; and a GUI for real-time manipulation of model parameters. Xolotl’s ease of use makes it an attractive option for pedagogical applications, rapid prototyping of models, and primary research use. Our software aims to simplify the investigation of the dynamics of conductance-based network and neuron models, facilitate collaborative modeling, and is intended to complement other tools being developed in the computational neuroscience community.

## 2 DESIGN GOALS

Xolotl is designed to be easy-to-use and richly featured while being fast enough to use in everyday research. Our focus was on designing an approachable simulator of conductance-based neurons and networks of these neurons; simulating arbitrary dynamical systems is therefore beyond the scope of this software. Specifically, the software was designed to simulate models of the form

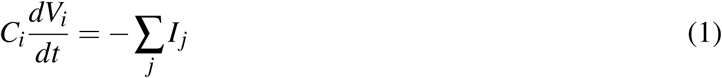

where *C_i_* and *V_i_* are the capacitance and membrane potential of compartment *i*. Compartments can represent whole neurons or parts of neurons. *I_j_* is the current due to ion channel population *j* and is given by

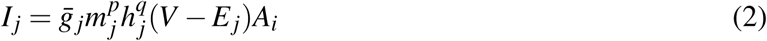

Here, 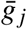 is the maximal conductance, and *E_j_* is the reversal potential of the ion channel population *j*. *A_i_* is the surface area of compartment *i* that contains these ion channels. *m_j_* and *h _j_* are activation and inactivation variables that change according to

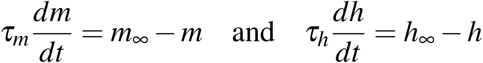

Typically, *τ_m_, τ_h_, m*_∞_, and *h*_∞_ are functions of the membrane potential *V_i_*. The software uses integration schemes that have been specifically developed to solve equations of this form (Dayan and Abbott 2001; Hines 1984; Oh and French 2006), though other schemes can be used if desired.

This software is designed to be used from within MATLAB, a scientific programming language common amongst neuroscientists and engineers for pedagogy and research. Our goal was to make xolotl completely usable entirely from within MATLAB. Models created using this simulator appear in the MATLAB workspace as native objects, are thus fully scriptable, and are fully compatible with the large library of toolboxes that MATLAB provides, allowing the software to be used as a component of other packages and tools. All parameters of a model, and activation functions of any channel can be inspected at any point. We designed several features of the simulator to be easily extensible: adding custom conductances or synapse types is possible by calling functions that generate new C++ files on-the-fly. Finally, xolotl is fully auditable by design, with several tools to verify model and parameter integrity and aid in reproducibility.

### 2.1 FEATURES

#### Object-oriented

Xolotl is designed to mirror the nested and hierarchical structure of networks and neurons. Biological neuronal networks are made up of neurons that are connected to each other with synapses. Each of these neurons contains within it a set of conductances and synaptic currents that contribute to its electrical behavior. Intracellular mechanisms can act within the cell, or parts of the cell, to modify and regulate conductances, synapses, or other dynamic properties of the cell. Similarly, a xolotl model can contain a set of compartments that can represent individual neurons. Each compartment can contain an arbitrary set of conductances. Compartments, conductances and synapses can contain mechanisms that can affect anything in the model. All objects in the xolotl model tree (compartments, conductances, etc.) are bonafide MATLAB objects with their own type and properties. This object-oriented programming paradigm naturally represents the hierarchical structure of biological networks that contain neurons that contain populations of ion channels, and makes constructing, integrating, and thinking about these models easier.

#### A rich library of network components

Xolotl comes packaged with hundreds of pre-existing components (compartments, synapses, conductances, and mechanisms) that can be used as building blocks to construct model neurons and networks. “Compartments” represent sections of membrane with a single membrane potential, intracellular Calcium concentration, and set of constituent components; and can represent either entire cells or parts of cells. Objects of type “conductance” represent populations of ion channels in a compartment that produce transmembrane currents. “synapse” objects connect two compartments together by introducing a current in the postsynaptic compartment that depends on the presynaptic compartment’s membrane potential. Objects of type “mechanism” can represent any intracellular mechanism and can read and modify any other component in the cell, and can run arbitrary dynamical systems within them. Parameters of any of these objects can be easily inspected and modified at any time, either manually or through a programmatic interface.

#### Automatic type system

To circumvent the tradeoff between high-performance but hard-to-use languages like C++ and richly-featured but potentially slow languages like MATLAB, we constructed an automatic type system that links object oriented code in C++ to object oriented code in MATLAB. This architecture lets us construct the core of the simulator in C++, leveraging features of C++ like pointers that are not readily available in MATLAB. A rudimentary way to make this C++ code useable in MATLAB would be to re-write that code in MATLAB so that MATLAB objects can be bound to their C++ implementations. However, this approach is cumbersome and inefficient, and can introduce errors. Instead, our automatic type system creates objects in MATLAB on-the-fly from C++ class specifications, obviating the need to rewrite code in MATLAB while preserving a tight coupling between objects in the MATLAB workspace and their C++ implementation. Crucially, this method makes developing new code much easier and simplifies the task of constructing complex frameworks that span these two languages.

#### Automatic hashing and compiling

Because every model requires a compiled binary to run, a potential stumbling block is the problem of unambiguously linking a model to a binary executable. Xolotl solves this problem by hashing (Rivest 1992) the C++ header files of every component in the model recursively, allowing a model, no matter how complex, to be compactly represented by a short alphanumeric identifier (its “hash”). Compiled binaries are named using this hash, ensuring both that the correct binary is run to integrate the model, and that compilation occurs only as needed. This powerful feature enables the user experience to remain entirely within the MATLAB workspace, with compilation and selection of the correct binary occurring silently in the background.

### 2.2 LIMITATIONS

Our focus on xolotl’s ease-of-use and speed imposed some limitations on its feature set.

#### Limited to conductance-based models

xolotl has been developed specifically for conductance-based models. It does not currently support rate- or current-based models, or arbitrary dynamical systems.

#### Limited numerical integration strategies

Most components in the software are integrated using the exponential Euler method, which has been used in integrating neuronal models (Oh and French 2006; Dayan and Abbott 2001). However, it may be desirable to use other methods under certain conditions. It is possible to introduce new components that implement other integration schemes, or to modify the integration schemes of existing components, but that requires writing new C++ code. Currently, xolotl can only implement integration schemes with fixed step size.

#### Inefficient tools for handling large networks

xolotl was designed to work with small but complex networks and models, where every compartment and component is named, rather than numbered. It is more therefore suited towards simulating small, heterogeneous networks rather than large, homogenous networks. While the software can integrate large networks (> 1000 compartments), other tools are presumably more suited to this task, offering more natural frameworks for dealing with a large number of identical units. Similarly, xolotl is not optimized to solve coupled ODEs on complex branched morphologies, that other simulators like NEURON (Hines and Carnevale 1997) are specialized for.

#### New mechanisms require new C++ code

Adding new network components requires writing new C++ code. A new conductance in the Hodgkin-Huxley formalism (Hodgkin and Huxley 1952b; Hodgkin, Huxley, and Katz 1952; Hodgkin and Huxley 1952a; Dayan and Abbott 2001) requires creating a new C++ header file, though this is generally trivial. Implementing a new integration scheme or component type requires much more in-depth knowledge of the underlying C++ core code.

## 3 USAGE EXAMPLES

In this section, we illustrate how xolotl can be used to generate, inspect, and simulate a variety of models. These examples have been chosen to demonstrate various features of xolotl, and are intended to serve as templates upon which researchers and educators can build.

### 3.1 SIMULATING A HODGKIN-HUXLEY MODEL

The axon of the giant squid contains a fast inactivating sodium conductance (`NaV`), a slower noninactivating potassium conductance (`Kd`), and a passive leak current (`Leak`). Seminal work by Hodgkin and Huxley showed that depolarizations of the membrane could lead to an activation of the voltagesensitive `NaV` channels, which led to a run away depolarization that was terminated by the inactivation of `NaV` channels and the activation of `Kd` channels that repolarized the membrane (Hodgkin and Huxley 1952b; Hodgkin, Huxley, and Katz 1952). As one of the simplest models of excitable neural membranes, the Hodgkin-Huxley model often serves as a the first model introduced in pedagogical literature (Dayan and Abbott 2001; Sterratt 2011; Trappenberg 2010).

In this example, we demonstrate how to simulate the spiking activity of a Hodgkin-Huxley-like model, and how the tools built into xolotl make it easy to set up and integrate the model and gain insight into the underlying biophysical mechanisms. This simple model consists of a single electrical compartment with three types of conductances (Figure 1A). This hierarchical organization of the neuron is mirrored in the structure of the model in the simulator: an object of type “compartment” is used to represent the cell body, and three objects of type “conductance” are used to model the three populations of ion channels (Figure 1B). These models of ion channels were obtained from (Liu et al. 1998) based on electrophysiological recordings from the lobster stomatogastric ganglion (Turrigiano, LeMasson, and Marder 1995), and are part of the simulator. The code to set up this model is terse, idiomatic, relies on no special markup, and preserves the hierarchical nature of the model (Figure 1C).

**Figure 1:**
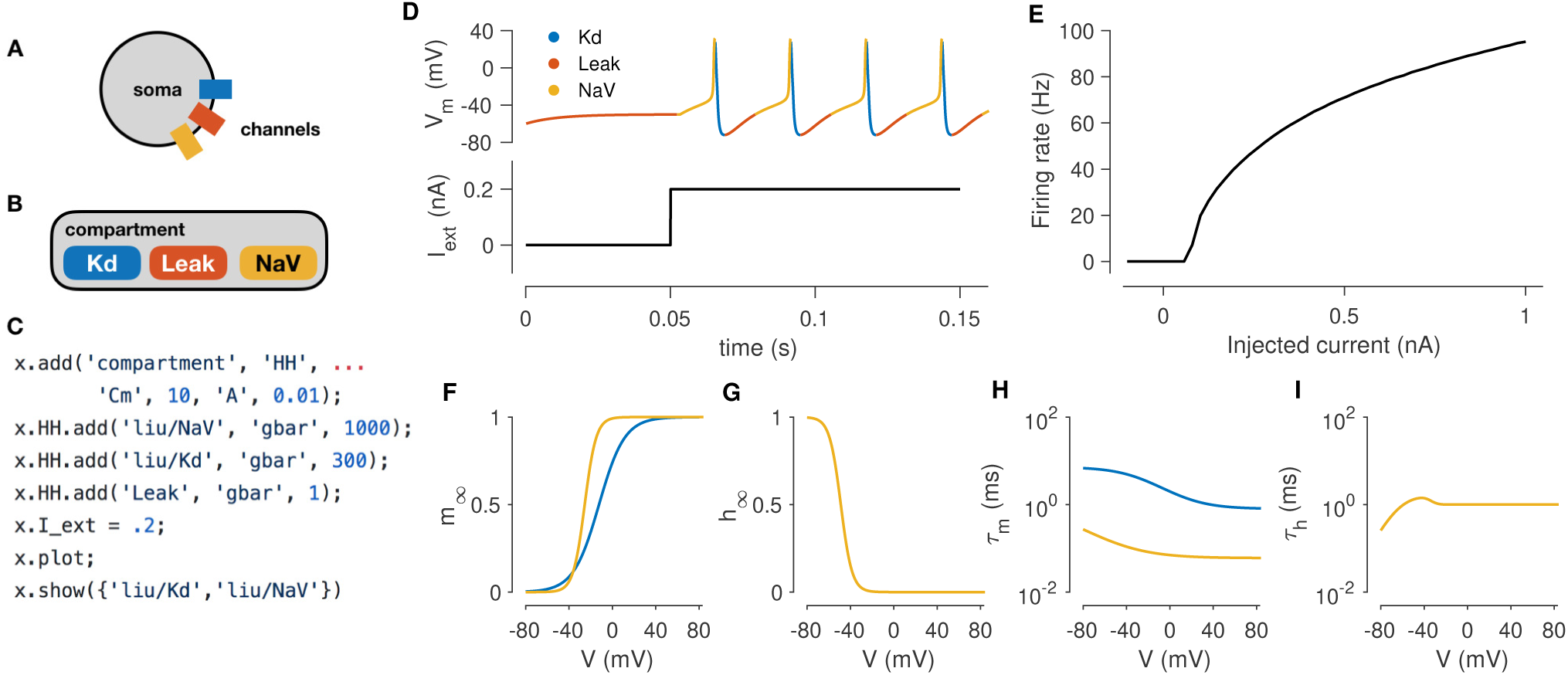
Simulating a single-compartment Hodgkin-Huxley spiking neuron model. (A) Schematic representation of a single-compartment neuron with three populations of ion channels (colored rectangles). (B) In xolotl, the soma is represented using an object of class “compartment” and populations of ion channels are represented by “conductance” objects contained within the compartment object. (C) The code snippet shown sets up this neuron model, injects current, integrates and plots the voltage, and displays activation functions, all in a few lines of code. (D) Simulated voltage trace of a Hodgkin-Huxley model with three conductances and 0.2 nA of injected current. Colors indicate the dominant current (gold is fast sodium (NaV), blue is delayed rectifier (Kd), red is Leak). (E) firing rate *vs*. current (f-I) curve of this neuron. (F-G) Steady-state gating functions for activation (m) and inactivation (h) gating variables. (H-I) Voltage-dependence of time constants for activation (m) and inactivation (h) gating variables

Adding an injected current and calling the built-in `plot` function plots the time series of membrane voltage. In the absence of injected current, the model is quiescent. When 0.2 nA is injected, the model tonically spikes (Figure 1A-B). The `plot` function displays a voltage trace colored by the dominant current (Figure 1A). Colors in the voltage trace indicate the strongest instantaneous inward current when the voltage is increasing, and strongest instantaneous outward current when the voltage is decreasing. This built-in feature reveals that the dominant current during the upswing of every action potential is the sodium current, and the dominant current immediately after the peak of the action potential is the potassium current, but that the leak current contributes to depolarization following an action potential. This feature could be useful in quickly understanding the contributions of a number of ion channel types in a complex voltage trace from a more complicated neuron model.

Integrating the model returns the voltage time series for every compartment:

~~~
V = x.integrate;
~~~

The model can be integrated for various amplitudes of injected current to determine its F-I (frequency current) curve (Kispersky, Caplan, and Marder 2012). Figure 1E shows the F-I curve of this model, obtained by repeated integration of the model. Finally, the built-in `show` function can plot activation (*m*) and inactivation (*h*) curves and voltage-dependent timescales of any channel type in the simulator (Figure 1D-G). These plots reveal that activation kinetics of the `NaV` channels are much faster than that of the `Kd` channels (Figure 1F), which facilitates the transient depolarization in an action potential. In summary, the simulator allows the user to construct and integrate this model in a few lines of code, and provides rich visualization of the dynamics of the model.

### 3.2 PERFORMING A VOLTAGE CLAMP EXPERIMENT *IN-SILICO*

Voltage clamping is an experimental technique where a amplifier is configured to inject the appropriate amount of current through a electrode to maintain the voltage of a cell at a desired value (Dayan and Abbott 2001). Under this paradigm, the membrane voltage is “clamped” or fixed to a desired value, permitting the study of voltage-dependent ion channels, since the sum of all currents through the population of ion channels in the cell is equal and opposite to the current injected by the clamp (Figure 2A). By combining voltage clamp with the use of pharmacological agents to block all channels but the one of interest, the voltage-sensitivity of an ion channel population can be characterized (Cole and Moore 1960; Cole 1955; Hodgkin and Katz 1949; Hodgkin, Huxley, and Katz 1952; Hodgkin and Huxley 1952a; Turrigiano, LeMasson, and Marder 1995).

**Figure 2:**
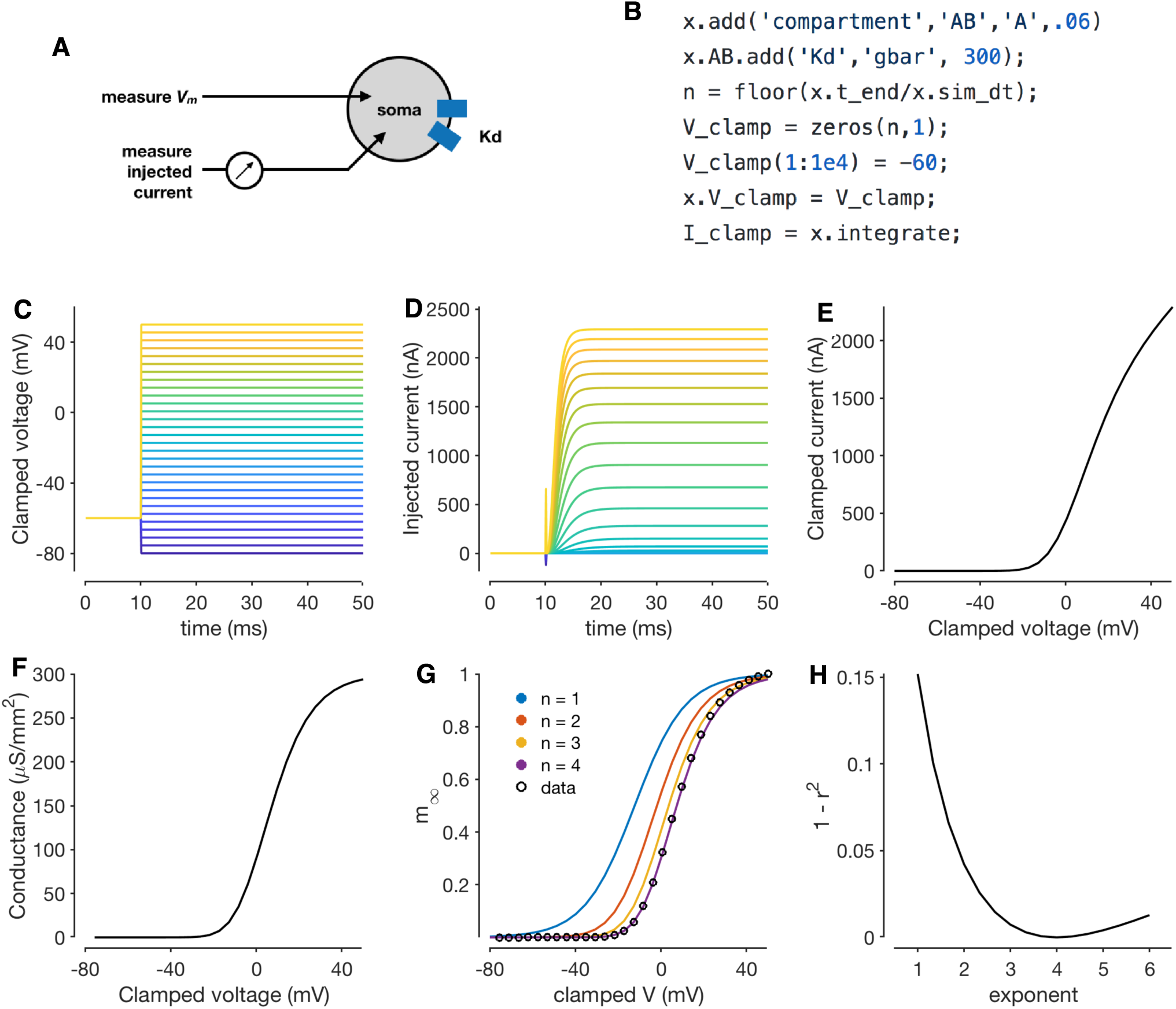
Simulating a voltage-clamp experiment. The diagram shows a cell with delayed rectifier potassium conductance (Liu et al. 1998) that is being recorded from in two-electrode voltage clamp (A). The code snippet shown here sets up a model with a single compartment and a single channel type, and clamps the cell to a constant voltage and integrates it (B). Voltage steps that the cell is clamped to (C). Clamp currents as a function of time (D). Asymptotic clamped current *vs.* clamped voltage for this cell (E). Accounting for the reversal potential of Potassium ions yields the conductance-voltage curve of this channel type (F). Normalized conductance-voltage curves, with sigmoid fits with various exponents (G). An exponent of *n* = 4 yields the best fit, allowing for the characterization of the activation function of this channel type (H).

Xolotl can reproduce such a voltage clamp experiment *in-silico*. Figure 2B illustrates how a simple model with a single compartment and a single ion channel type can be set up and clamped to a desired voltage. Integrating the model yields the current required to clamp the cell at that voltage. Here, we use a delayed-rectifier potassium conductance (Liu et al. 1998) and simulate a voltage-clamp experiment whose goal is to infer the activation function of this channel. First, the cell is clamped to a number of different voltages (Fig. 2C) and the resultant clamp currents are measured by integrating the model (Fig. 2D). Since the compartment is being voltage clamped, integrating the model returns the clamping current:

~~~
I_clamp = x.integrate;
~~~

By repeating the integration at a number of clamp voltages, we observe that the asymptotic clamp currents depend on the clamp voltage in a nonlinear manner (Fig. 2E), since the open probabilities of the channel are functions of the membrane voltage. Assuming the reversal potential is known, Eq. (2) can be used to solve for the total conductance of the channel as a function of the clamp voltage (Fig. 2F). Finally, a sigmoid can be fit to the normalized conductance-voltage curves to obtain the activation function of the ion channel population (Fig. 2G-H). Xolotl can therefore be used to describe graphically the theoretical underpinnings of ion channel characterization through voltage clamp and can serve as an effective pedagogical tool in computational neuroscience.

### 3.3 INTRACELLULAR MECHANISMS

So far, the models we described only considered the voltage dynamics of a cell (the solution to Eq. 1). However, real neurons possess several dynamical features, arising from a variety of intracellular mechanisms. Xolotl makes it possible to model and include arbitrary intracellular mechanisms. In xolotl, these intracellular mechanisms are represented by the “mechanism” object, and can be bound to compartments, conductances, and other object types.

A key intracellular mechanism is the cytosolic buffering of Calcium and its influx through voltagegated Calcium channels. Figure 3A shows a model of a single-compartment model with 8 populations of ion channels (Liu et al. 1998). Without any explicit mechanism for Calcium influx or buffering, the intracellular Calcium levels in this model do not change (Figure 3B) and the model tonically spikes (Figure 3C). Calcium buffering and influx can be modeled by a differential equation that increases intracellular Calcium with the current through Calcium channels and relaxes back to a baseline value (Liu et al. 1998; Prinz, Billimoria, and Marder 2003; Dayan and Abbott 2001) (Figure 3D). This mechanism exists in the xolotl library as `CalciumMech1` and can be added to the model using a simple `add` statement (Figure 3E). The intracellular Calcium in the model now oscillates periodically (Figure 3F), synchronized to bursts in action potentials in this cell (Figure 3G).

**Figure 3:**
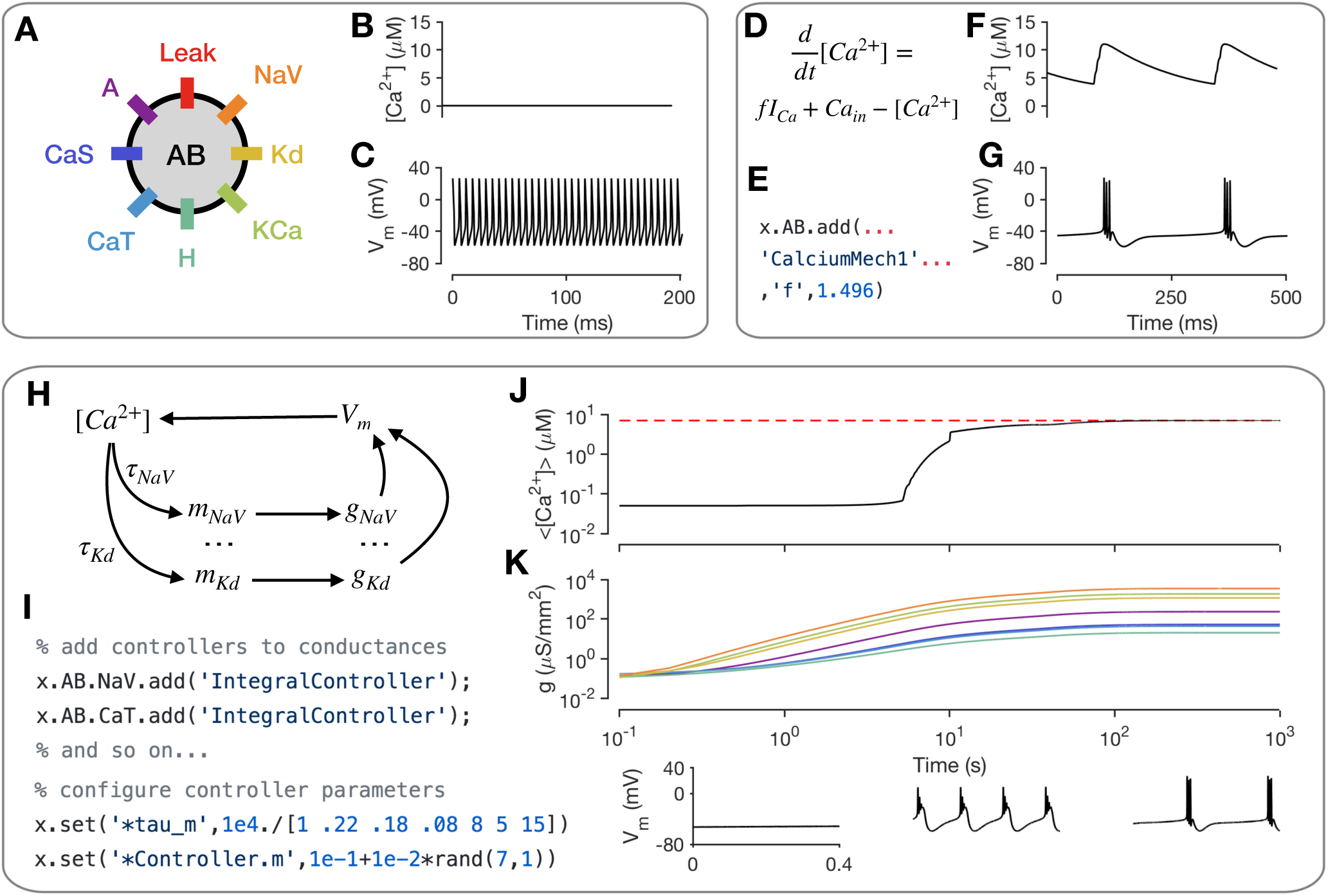
Modeling intracellular mechanisms. A single-compartment neuron model with 8 channel types (A). Because there is no mechanism for changing intracellular Calcium in this model, the Calcium level stays constant (B), and the cell tonically spikes (C). Intracellular Calcium buffering and influx through voltage gated Calcium channels (VGCCs) can be modeled using a simple differential equation (D). Code snippet shows how this mechanism can be added to the neuron model (E). The cell now bursts periodically, with synchronized oscillations in intracellular Calcium (F-G). Schematic of Calcium-dependent integral control homeostasis (O’Leary et al. 2013; O’Leary et al. 2014) (H). In this feedback system, the rates of mRNA synthesis depend on the Calcium level in the cell, which depends on the membrane voltage, which in turn depends on the conductance density of all channel types, which, through translation, depends on the mRNA abundance. (I) The code snippet shows how these integral controllers are implemented as mechanism objects, and can be added to conductances. (J) On integrating the model, intracellular calcium levels rise and approach the target (red dashed line). This is accompanied by an increase in the conductance densities of all channels being controlled by this homeostatic mechanism (K). The voltage behavior of the cell changes from silence to bursting with truncated spikes to regular bursting.

Neurons can regulate their electrical activity by controlling the spectrum of ion channels they express (MacLean et al. 2003; Turrigiano, LeMasson, and Marder 1995; Schulz, Goaillard, and Marder 2006). Here, we will show how xolotl can be used to represent a recently proposed model of a homeostatic feedback system that controls the transcription rates of ion channels with the integral of an error signal derived from the intracellular Calcium concentration (O’Leary et al. 2013; O’Leary et al. 2014) (Figure 3H). Since this mechanism affects each ion channel population individually, an object corresponding to this mechanism is added to each conductance object in the neuron (Figure 3I). Setting all maximal conductance densities to some low value and integrating the model shows that the intracellular Calcium levels rise over time and approach the target Calcium concentration (Figure 3J), while all conductance densities increase and then remain bounded (Figure 3K). Examining the voltage dynamics of the cell reveal that it transitions from quiescence to truncated bursts of action potentials to periodic bursting as this mechanism regulates the neuron’s ion channel spectrum. In summary, xolotl can be used to construct neuron models with intracellular mechanisms such as Calcium influx and buffering, and homeostatic regulation.

### 3.4 USING SNAPSHOTS TO EXPLORE MODEL DYNAMICS AND PARAMETERS

Switching back and forth points in parameter space and state space of a neuron model is a common occurrence in working with neuron models, and a significant fraction of a modeler’s time is spent in a feedback loop of running simulations, viewing the output, changing parameters, and re-running simulations (De Schutter 1992). Xolotl makes it easy to bookmark configurations of a model and return to them at will. Internally, xolotl uses the `serialize` method to gather all parameters and dynamic variables into a vector of values that is passed to the underlying C++ implementation. A paired `deserialize` method is used to update all parameters and variables in the object tree from this vector. This architecture provides a natural framework for representing the state of any model, no matter how complex, using a vector of numbers. The `snapshot` method built into xolotl leverages this schema to save the entire state of the model in a named variable, that can be accessed using another built-in method called `reset`.

(Figure 4) illustrates how these features can be used to explore the dynamics of the model presented in the previous section. First, the current state of the model is saved using the `snapshot` method into a state called “initial”. In this state, the model exhibits periodic bursting activity due to a particular configuration of maximal conductance densities (Figure 3, orange). On setting the maximal conductances of the Calcium-permissive channels to 0, the model switches to a tonic spiking activity (Figure 3, purple). Integrating the model for a longer duration allows the homeostatic control mechanism in the cell to restore the conductance profile and bursting activity to a state close to the original state (Figure 3, green). This state is saved using the name “final”.

**Figure 4:**
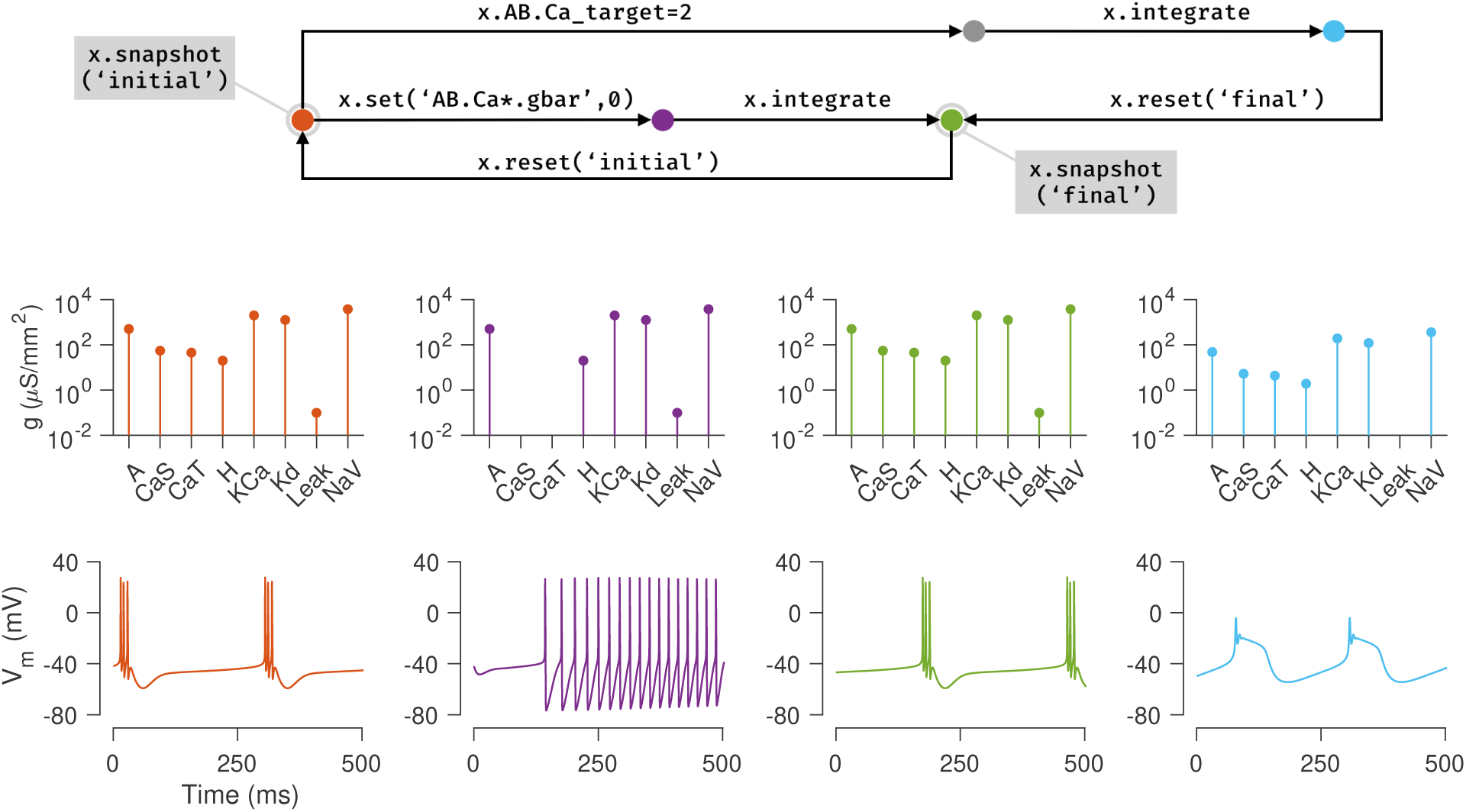
Snapshots allow the user to bookmark points in parameter and state space of the model and return to them *ad arbitrium*. The initial state (orange node) of the single compartment model in the previous example is saved using the `snapshot` method. This method saves all parameters and dynamic variables of the model in a named state. The first column (orange plots) shows the profile of conductances and the voltage dynamics of the model at this point at this time. The maximal conductances of the Calcium channels are then set to zero (purple node), changing the voltage dynamics of the neuron (purple plots). After evolving the model for some time (green node), the conductance profile and voltage dynamics returns to a state similar to the initial state (green plots). This configuration is now saved in a state called *f inal* and the initial configuration is returned to using the `reset` method (backwards arrow from green to orange). Another parameter is now changed (the Calcium target), and the model is integrated to reach a new state (blue node) where the voltage dynamics are different from the *initial* state. In summary, any state can be bookmarked using a descriptive name using the `snapshot` method, and can be returned to using the `reset` method.

The initial state can be returned to using the `reset` method, and a new manipulation to the model can be explored. Here, the intracellular Calcium target is modified, and the model is re-integrated, to yield a different voltage activity (Figure 3, blue). At the end of this numerical exploration, any of the saved states can be quickly returned to using the `reset` method, making the process of re-initializing models to desired states and parameters both error-free and efficient.

### 3.5 SIMULATING NETWORK MODELS

Neurons communicate and interact using synapses, electrochemical junctions between cells (Gjorgjieva, Drion, and Marder 2016; Hua and Smith 2004). In chemical synapses, the presynaptic neuron releases packets of neurotransmitter across the synaptic cleft, which activate receptors on the postsynaptic neuron. In electrical synapses, no chemical intermediary is involved. New patterns of activity can emerge from the characteristics of the connecting synapses (Li, Bucher, and Nadim 2018; Nadim et al. 1999; Gutierrez and Marder 2013; Gutierrez, O’Leary, and Marder 2013).

Network models in xolotl consist of compartment objects that can be connected by synapse objects. Two compartments representing different neurons can be connected using synapses using the built-in `connect` method. For example, to connect two single-compartment neurons called `LP` and `PY` using a chemical synapse of type `Cholinergic` with strength 30 nS, we can use

~~~
x.connect(‘LP’, ‘PY’, ‘Cholinergic’, ‘gbar’, 30);
~~~

Xolotl has several types of synapses built-in, and other synapse classes can be easily added using templates. Figure 5 demonstrates the implementation of a model of the triphasic pyloric rhythm in the stomatogastric ganglion of crustaceans (Prinz, Bucher, and Marder 2004). The pyloric model contains three compartments (`AB, LP`, and `PY`) and seven synapses (Figure 5A). This structure is recapitulated in the hierarchy of the xolotl object, where conductances are contained within compartments (Figure 5B). The membrane potentials show triphasic rhythmicity in the three compartments (Figure 5C-E). When the `PY` is depolarized, the dynamical variable mediating the glutamatergic (`Glut`) synapse model between `PY` and `LP` is close to 1 and `LP` is inhibited (Figure 5F). Conversely, when `PY` is hyperpolarized, the dynamical variable is close to 0. In this model, when `PY` spikes, IPSPs can be seen in the `LP` voltage trace (Figure 5D-E).

**Figure 5:**
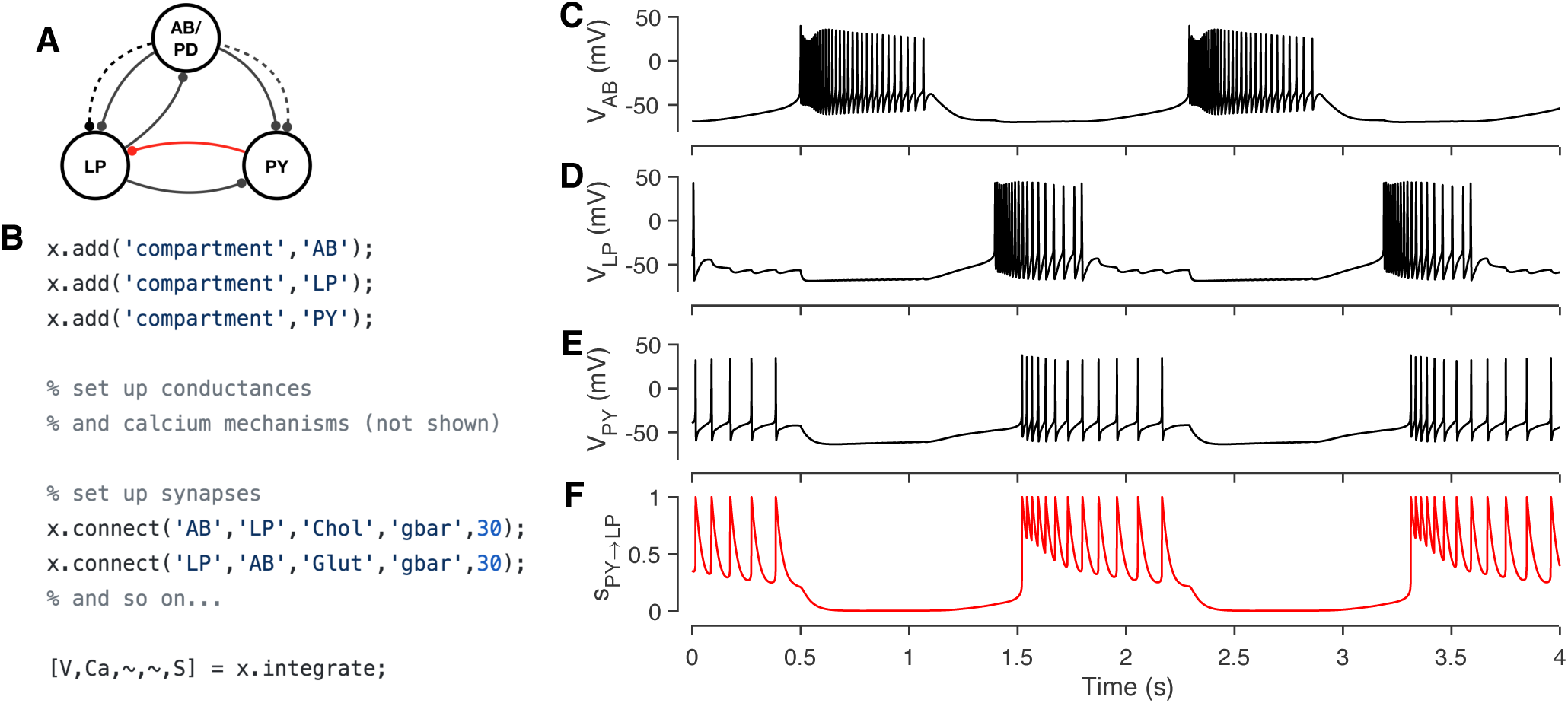
Simulating a network of conductance-based model neurons, coupled by voltage-dependent dynamic chemical synapses. A three-compartment model of the pyloric network in the crustacean stomatogastric ganglion (Prinz, Bucher, and Marder 2004) (A). Each neuron is modeled with a single compartment with 7-8 intrinsic conductances and 1-3 post-synaptic conductances. Synapses can be one of two types (Cholinergic (dashed lines) or Glutamatergic (solid lines)) and have different kinetics. The code snippet shows how neurons can be created, wired together using synapses, and how the model can be integrated to return voltages and intracellular calcium levels in every compartment, and the state of every synapse (B). Simulated voltage trace of a model network for the three compartments obtained from this simulation (C-E). Time series activation variable of the Glutamatergic synapse between PY and LP (red connection in diagram) shows how the synapse becomes active every time the PY neuron spikes (F).

### 3.6 USING THE GUI TO MANIPULATE PARAMETERS

Conductance-based neuron models are typically high dimensional and contain many parameters. Changing a single parameter monotonically can cause non-monotonic changes in behavior of the model, and certain dynamical features may only emerge when in specific non-convex regions of parameter space (Golowasch et al. 2002). It is often challenging to build intuition about what effect a parameter has on the model under these conditions. Traditionally, the technique used by computational neuroscientists in building intuition about these models is to iteratively run simulations, view outputs and change parameters. In practice, this meant writing a script, running it, inspecting the output, changing parameters in the script, and repeating this process. It can be cumbersome, and every step in this process involves “mode” changes: switching between a text editor, viewing a graphical output, and the command line that can frustrate the researcher.

Xolotl is designed to streamline this process and allows for any parameter in any model to be manipulated using graphical sliders, with immediate, real-time feedback of its behavior. Any model in the simulator can be manipulated using

~~~
x.manipulate
~~~

This method creates a GUI element with sliders for every parameter, and also creates a set of plots that shows the dynamical behavior of the model (Fig. 6). By default, time series of the voltage and the calcium of every compartment are shown, though this can be modified. Moving any of the sliders updates the value of that parameter in the model, and also triggers a function call that reintegrates the model and updates the output plots. This function call can also update custom plots, like the one shown in (Fig. 6B). Any model can be manipulated in this way without writing any additional code.

**Figure 6:**
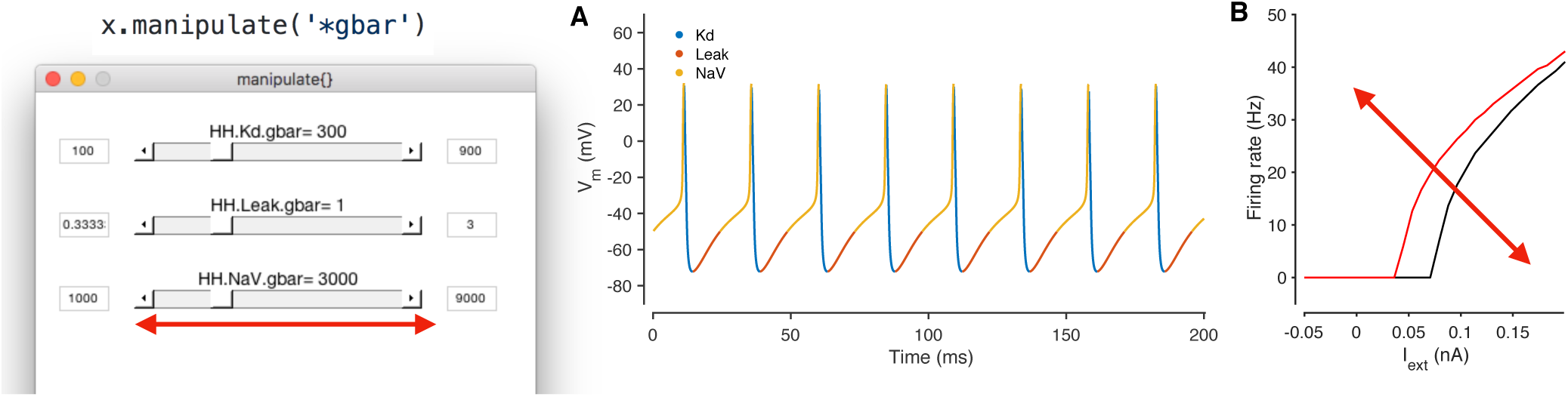
Manipulating neuron parameters in real time. Any set of parameter in the model can be manipulated; here, the maximal conductance of every conductance type in the model from Fig. 1 is being manipulated using the code snippet shown here. The screenshot shows a GUI with sliders for every parameter of interest that is created by the `manipulate` method. These sliders can be linked to an arbitrary number of visualization functions. In this example, two visualization functions are used: the built in `plot` method (A) and a custom function that computes the firing-rate-*vs*.-injected current curve for this neuron (B). Both plots refresh themselves with every movement of any slider, allowing the user to build intuition about how every parameter controls the dynamical behavior of the model. A screen recording of this model being manipulated in real time is included in Supplementary Material

This feature was only possible due to our architectural decision to split the code base across two programming environments. A rich scientific programming environment like MATLAB makes it possible to easily generate user interface elements and to bind them to data in plots, while the sheer speed of compiled languages like C++ allow for the immediate, real-time feedback and updating of plots. By default, any parameter in the model can be manipulated, including parameters in user-defined mechanisms that do not exist in the base simulator. The GUI can be constrained to arbitrary subsets of model parameters using wild card matching (as shown in Fig. 6) or by manually specifying the parameters of interest.

## 4 BENCHMARKS

Our goal in designing xolotl was to create an easy-to-use neuron and network simulator that was fast enough and accurate enough for routine use by computational neuroscientists. In this section, we benchmarked the speed and accuracy of xolotl in simulating single and large numbers of Hodgkin-Huxley-like neuron models (as in Fig. 1) and bursting neuron models based on the bursting neurons in the lobster stomatogastric ganglion (STG) (Prinz, Bucher, and Marder 2004). For each type of neuron model, we compared our software to NEURON (Hines and Carnevale 1997), a high-performance and powerful neuron simulator specialized in simulating neurons with complex morphologies and DynaSim (Sherfey et al. 2018), a general-purpose simulator that can solve coupled differential equations numerically. All simulators were run on the same hardware using fixed time-step solvers: xolotl used the Exponential Euler method (Dayan and Abbott 2001), NEURON used the implicit Euler solver (Hines and Carnevale 1997) and DynaSim used C-compiled 2^nd^-order Runge-Kutta integration scheme as recommended for high-performance (Sherfey et al. 2018). We measured the speed of each simulator by dividing the time simulated for, by the time it took for the simulator to complete integration. For example, if a simulator could simulate 10 seconds of model data in 1 second, its speed would be 10X. We measured the speed of every simulator as a function of simulation time step, total length of simulation, and system size.

All three simulators were faster with larger time steps, since fewer iterations were needed (Fig. 7A-B), and were approximately linear in the region tested. Xolotl compared favorably to NEURON and DynaSim in this task. We also measured the quality of the simulated output by comparing it to the simulated output at the smallest time step. Simulation error was measured using the LeMasson cost (LeMasson and Maex 2000), and was comparable amongst the three simulators (Fig. 7C-D). Since xolotl sets up and runs the simulation in C++, it needs to transfer parameters and data to and from the underlying implementation. To measure the performance cost of this overhead, we repeated these benchmarks on all three simulators at a fixed time step of 0.1 ms and varied the length of time simulated for. Speed increased with simulation duration up to a point, and then saturated, indicating a fixed performance cost to the overhead (Figure 7E-F, black lines). Simulations using DynaSim, which also used a similar architecture and need to move data between C implementations and the MATLAB workspace, showed a similar increase in speed with simulation length (Figure 7E-F, red lines). However, simulations using NEURON ran at a constant speed irrespective of simulation length, presumably due to differences in the underlying implementation (Figure 7E-F, blue lines).

**Figure 7:**
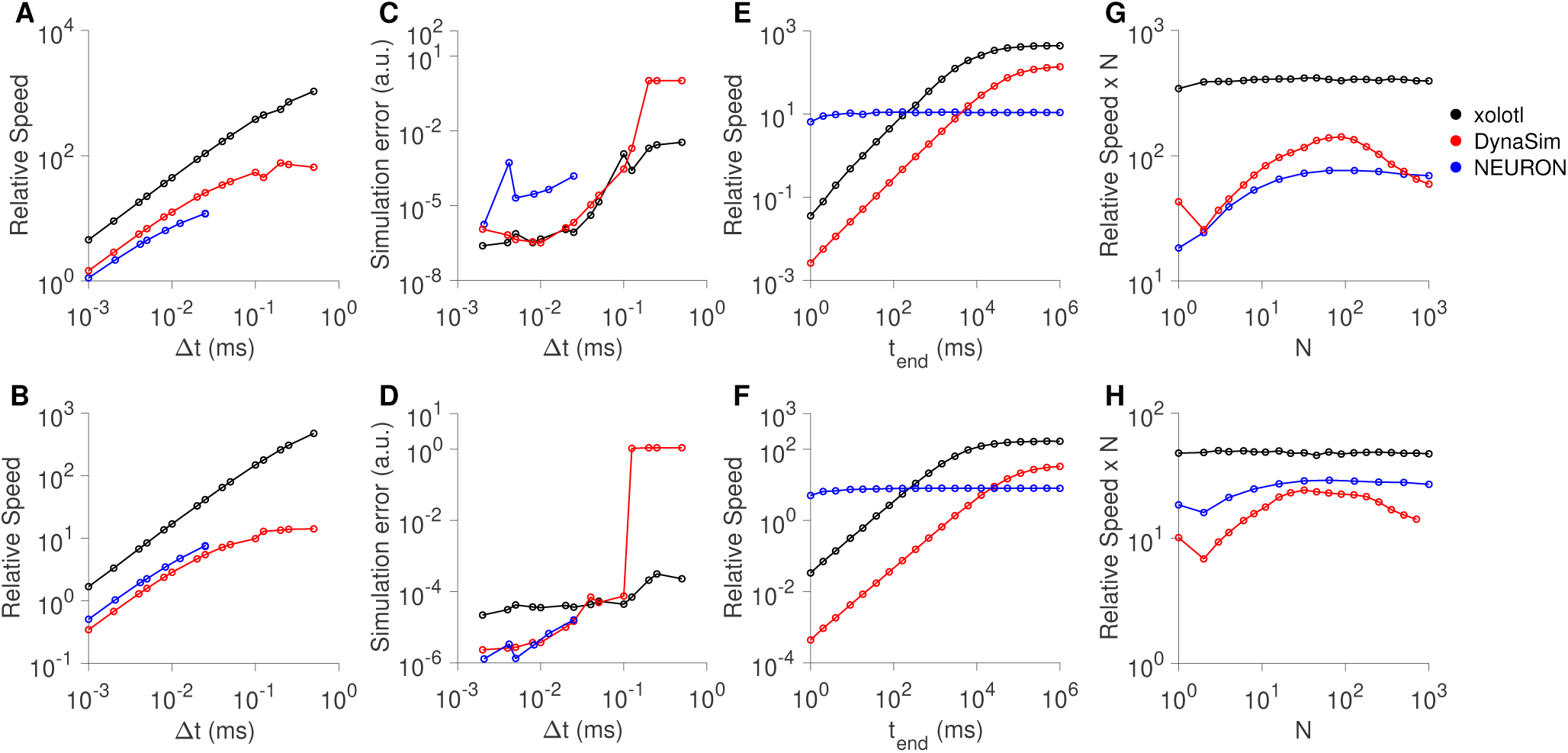
Comparison of speed and accuracy of xolotl, NEURON and DynaSim. The top row shows simulations of a tonically-firing Hodgkin-Huxley model with three conductances with constant injected current (as in Fig. 1). The bottom row shows simulations of a bursting stomatogastric ganglion neuron model with 8 conductances (as in Fig. 5). Ratio of run-time to simulation time (relative speed) as a function of simulation time step (A, B). Simulation error as a function of the step size (C, D). Relative speed of integration as a function of the simulation length (E, F). Relative speed of integration, normalized by system size, as a function of the number of compartments simulated simultaneously (G, H). All benchmarks were performed on the same computer, and all simulators using fixed time-step integration methods. NEURON was run using the Python wrapper.

Many simulators have been designed with a focus on simulate large numbers of compartments, either as networks with many identical neurons or in a large multi-compartment neuron model (Brette et al. 2007; Sherfey et al. 2018; Vitay, Dinkelbach, and Hamker 2015; Delorme and Thorpe 2003). While our software is not designed for this task *per se*, we measured its performance as a function of the number of compartments simulated. Xolotl can quickly create a number of identical copies of a compartment using the `replicate` method:

~~~
x.replicate(‘compartment_to_replicate’,n_copies);
~~~

We used the `replicate` method to create and run models with varying numbers of neurons (either Hodgkin-Huxley-like or bursting neurons) and measured the speed of all three simulators as a function of system size. Plotting the speed normalized by the system size *vs.* the system size, we observed that the speed of integration of xolotl is linear with system size (Figure 7G-H), black lines), for up to 1000 single-compartment neurons (up to 13,000 ODEs). Its performance compares favorably with that of NEURON and DynaSim as a function of system size (Figure 7G-H).

## 5 DISCUSSION

We set out to design a neuron and network simulator that could be useful in the classroom setting, especially for students of computational neuroscience, while also being powerful, fast, and extensible enough to be used for research. By using a novel architecture that permits the symbolic manipulation of C++ objects in a intuitive MATLAB interface, we demonstrated some of the features of xolotl using simulations of single-compartment models of Hodgkin-Huxley like neurons (Fig. 1); voltage clamp experiments to recover activation functions of single channel types (Fig. 2); a neuron model where intracellular mechanisms can control the dynamics of Calcium and can regulate the maximal conductance of ion channel types in an activity-dependent manner (Fig. (3); and a network of neurons with multiple synapse types (Fig. 5). We also illustrated how built-in features of the simulator make it easy to bookmark and jump between model configurations (Fig. 3, purple), and how parameters of the model can be changed using sliders and their effect can be viewed in real time (Fig. 6).

### 5.1 A FOCUS ON USABILITY

“About half the time spent on a typical simulation project involves creating and tuning the model. Thus, a good user interface may contribute more to the overall efficiency of a project than pure computation speed.” (De Schutter 1992). Xolotl is designed primarily with ease-of-use in mind. This includes the time it takes to install, setup, and learn how to use the software, the time to write and debug scripts, and the time to perform the simulations (Rudolph and Destexhe 2007). An easy-to-use simulation environment must minimize time spent in all these domains, especially during human engagement with the software. Complicated software remains broadly inaccessible and time-consuming even to perform single-compartment simulations, though the actual simulation time may be very small.

We have focussed on making our software as easy to use as possible, without sacrificing performance or extensibility. For example, the software and all dependencies can be installed using a single-line installer script from within the MATLAB command line. The installation includes worked example scripts that demonstrate various features of the simulator, that can be run without any configuration. We decided to built the the front-end interface to xolotl in MATLAB to facilitate interoperability with existing tools for time series analysis, optimization, and parallel computing. This allowed us to build rich tools for visualization and to interact with the simulation.

### 5.2 COMPARISON WITH OTHER SIMULATORS

Over the years, several simulators have been developed to integrate systems of coupled differential equations that model the spiking activity of neurons (Brette et al. 2007; Sherfey et al. 2018; Vitay, Dinkelbach, and Hamker 2015; Delorme and Thorpe 2003; Hines and Carnevale 1997; Bower, Beeman, and Hucka 2003). A critical architectural choice in designing a simulator is how much “scaffolding” a user is provided with to construct a model; whether a model is specified by equations or by components, or by some combination of the two. In an equation-oriented architecture, the user starts with a blank slate and the primary method of specifying a model is to write out its differential equations (Stimberg et al. 2014b). In contrast, in a component-oriented architecture, the primary method of specifying a model is to assemble it from pre-existing components, each of which include differential equations, parameters, and solvers. Both approaches have advantages and disadvantages that are discussed below.

An equation-oriented simulator can be more transparent and allows the user to know exactly what is being solved, but equations can be cumbersome to write out, read, or to debug. In most commonly used programming environments, these equations have to be entered as strings, and complex parsing has to be carried out by the simulator to check that these strings constitute valid equations. In addition, parameters have to be written explicitly into equations, and it is usually not trivial to change parameters after initial specification. In contrast, components are easy to assemble into a model, but it can be hard to know what they contain, where they are physically located on the user’s computer, and how they can be changed. NEURON is primarily a component-oriented neuronal simulator, and new components are specified in special model files that can be “inserted” into a model. BRIAN is an equation-oriented neuronal simulator meant to be used from within Python (Goodman and Brette 2009). XPP is a general purpose dynamical system simulator that is equation-oriented (Ermentrout 2002). DynaSim is an equation-oriented simulator with some component-oriented capabilities. Models can be specified by both strings of equations or components, but since models do not exist as objects in the workspace, parameters and variables have to be reinitialized when changed (Sherfey et al. 2018).

Xolotl is a purely component-based simulator, and all equations need to be included in a C++ header file that specifies a object. Since our automatic type system binds MATLAB objects to the underlying C++ header files, objects can be inspected and parameters can be modified in the MATLAB workspace. To mitigate some of the drawbacks of the component-oriented paradigm, we have implemented an architecture that allows the user to access the underlying C++ code of any object by simply clicking on the object tree in the MATLAB command line. This feature removes the uncertainty inherent in other component-oriented simulators of the equations underlying each component, and allows the user to modify these equations if needed.

Another architectural choice in designing simulators is the syntax required for specifying models. Equation-oriented simulators specify models as strings of equations, so must invent a new syntax to specify derivatives, variables, and other common elements in coupled differential equations. Some component-oriented simulators like NEURON also specify their own syntax, or have invented their own language to specify components and write out equations. While this allows for powerful features such as support for units in NEURON, the user is required to learn a novel syntax and vocabulary, hindering ease of use. In general, the syntax for model specification in different simulators can be different. For example, DynaSim, BRIAN, XPP, and NEURON all use different, incompatible formalisms to represent equations, increasing the cognitive load on users using more than one simulator. Here, we have elected not to specify a domain-specific “middleware” layer, and instead specify and implement models and equations in idiomatic C++. This greatly decreases the learning curve and allows users with a general familiarity with programming languages to quickly acquaint themselves even with the most technical parts of the simulator.

### 5.3 AUDITABILITY AND REPRODUCIBILITY

The use of computational tools is increasingly central to the scientific method; yet, the lack of auditability and accountability in their use has led to a crisis of credibility affecting many scientific fields (Stodden et al. 2016; Baker 2016). Unlike in experimental research, where a lack of reproducibility can manifest due to meaningful reasons like uncharted differences in experimental protocols or intrinsic variability, the reasons for irreproducibility in computational research are often trivial and include: a) typographical errors from transcribing model parameters and equations, b) obscure software design that leads to users not knowing precisely what equations the software is solving, or what parameters it is using, c) incompatibility with version control systems and rolling software development that leads to ambiguity in which the version of the software that was used to generate a particular result is unclear, and d) convoluted architectures that make it too complex for non-experts to understand the inner workings of the software (Xu, Xu, and Deng 2017; Sedano 2016; Vikström 2009).

Our software was designed with this threat model in mind, and has features that allow the user to answer the following questions in the affirmative: “Can I be sure that I am doing what I think I am doing?” and “Is it possible for others to reproduce *exactly* what I have just done?”. Our goal was to design software that would allow the user to verify for herself that the software was running as she intended, and to be able to reproduce results from others quickly and unambiguously. The primary design choice in our software that enables auditability and reproducibility is that every simulation is tied to an alphanumeric checksum, or hash, using the MD5 algorithm (Rivest 1992). The hash is computed from every C++ file that is part of the model, and any changes in any C++ file included in the model will trigger a recompilation of that model. Thus, the hash guarantees that a given model is derived from a set of source files, obviating any ambiguity about the code used to generate a model. In addition, every parameter and state variable in the model can also be hashed together with underlying code, allowing the user to generate a short checksum that guarantees with high probability that every aspect of the model – code, parameters, and initial conditions – are exactly as they should be.

Typographical errors from transcribing model parameters and equations can also be detected using hashes, and the component-oriented architecture of our software makes it easy to debug code and spot errors. Our software has been designed so that it is possible to explore the model interactively in the command line, and it is possible to “click through” from the highest level of the model in the command line all the way down to the underlying code of any component in the model. This design allows the user to know precisely the equations being solved, and view the code that numerically integrates them. Finally, the core of our software is written in a few hundred lines of code and contains just four classes: compartments, conductances, synapses, and mechanisms. This architectural simplicity lets a motivated user understand the entirety of our code quickly.

### 5.4 OUTLOOK AND FUTURE DIRECTIONS

In its current form, xolotl is an efficient and easy to use neuron and network simulator that is actively being used in research. Results from an early version of this simulator guided intuition in the modeling of a recently characterized Calcium-dependent Potassium channel found in *Drosophila* neuromuscular presynaptic terminals (Bronk et al. 2018). Work on the simulator continues in the open at a publicly accessible repository (https://github.com/sg-s/xolotl/), and the library of conductances, synapses and mechanisms that xolotl ships with grows continuously. While this simulator was intended as a research tool, the many worked examples that are built into it demonstrate how it could also be used as a teaching tool.

Because xolotl models are bonafide MATLAB objects, they are compatible with most of the powerful tools that exist within MATLAB. For example, it is possible to write simple scripts that run xolotl models in parallel using MATLAB’s parallel processing toolbox, speeding up large simulations. Xolotl models are also compatible with the global optimization toolbox in MATLAB, allowing parameters in xolotl models to be optimized, enabling the creation of toolboxes that efficiently tune parameters in neuron models to satisfy arbitrary constraints (Achard and De Schutter 2006; Krichmar 2014; Keren, Peled, and Korngreen 2005; Van Geit 2007; Druckmann et al. 2008).

Care has been taken to reduce the amount of technical debt (Suryanarayana, Samarthyam, and Sharma 2014) associated with this project, with all parts of the simulator and dependencies written in a modular, objected oriented fashion. As a result, many of the key features and architectures of xolotl can be reused by others in their own applications. For example, cpplab, the automatic type system that binds C++ code to MATLAB objects, is independent of this simulator, and exists as a distinct repository that has been made freely available (https://github.com/sg-s/cpplab); and the framework for generating a GUI with sliders for each parameter that is hooked up to function callbacks also exists as an independent repository (https://github.com/sg-s/puppeteer) that can be easily integrated into other applications.

## AUTHOR CONTRIBUTIONS

SG-S & AH wrote the software and created the online user documentation. EM provided general oversight. All authors wrote and reviewed the paper.

## ACKNOWLEDGMENTS

The authors would like to thank Timothy O’Leary and Leandro Alonso for useful discussions, Janis Li for help transcribing many conductance models, and members of the Marder Lab, especially Sonal Kedia, Ekaterina Morozova, Jason Pipkin and Philipp Rosenbaum for careful reading of the manuscript.

## DATA AVAILABILITY STATEMENT

The code to generate all figures is available at (https://github.com/marderlab/xolotl-paper). xolotl is freely available at (https://github.com/sg-s/xolotl).

